# A cloud-based platform for the analysis of single cell RNA sequencing data

**DOI:** 10.1101/2020.09.28.317719

**Authors:** Nithin Joshy, Kyuson Yun

## Abstract

**Motivation:** Single-cell RNA sequencing (scRNA-seq) is a recent technology that has provided many valuable biological insights. Notable uses include identifying novel cell-types, measuring the cellular response to treatment, and tracking trajectories of distinct cell lineages in time. The raw data generated in this process typically amounts to hundreds of millions of sequencing reads and requires substantial computational infrastructure for downstream analysis, a major hurdle for a biological research lab. Fortunately, the preprocessing step that converts this huge sequence data into manageable cell-specific expression profiles is standardized and can be performed in the cloud. We demonstrate how a cloud-based computational framework can be used to transform the raw data into biologically interpretable cell-type-specific information, using either 3’ or 5’ transcriptome libraries from 10x Genomics. The processed data which is an order of magnitude smaller in size can be easily downloaded to a laptop for customized analysis to gain deeper biological insights.

**Results:** We produced an automated and easily extensible pipeline in the cloud for the analysis of single-cell RNA-seq data which provides a convenient method to handle post-processing of scRNA sequencing using next generation sequencing platforms. The basic step provides the transformation of the scRNA-seq data to cell-type-specific expression profiles and computes the quality control metrics for the dataset. The extensibility of the platform is demonstrated by adding a doublet-removal algorithm and recomputing the clustering of the cells. Any additional computational steps that take a cell-type expression counts matrix as input can be easily added to this framework with minimal effort.

**Availability:** The framework and its documentation for installation is available at the Github repository http://github.com/nj3252/CB-Source/

**Supplementary information:** Supplementary data available at *Bioinformatics* online.

## Introduction

Single-cell RNA sequencing is used to precisely determine expressed genes, and by extension, cell types and states among heterogenous cell types present within a sample (1). Application of this technology is rapidly growing and some of these are listed below to illustrate its power. Comparing the differences in gene expression between individual cells has identified novel cell-types in a tissue (2). In cancer research, the ability to find and characterize outlier cells has potential implications for furthering our understanding of drug resistance mechanisms and relapse in cancer treatment (3). scRNA-seq is also being utilized to delineate cell lineage relationships in early development, cell-type differentiation, and cell fate determination (4).

As with many genomics approaches, a major hurdle in using this technology is the amount of raw data that it generates and computational power needed to analyze the data. Higher-end computational infrastructure is required to process the raw sequencing data into biologically meaningful cell-type-specific expression profiles (5). However, the need for high power computational infrastructure is limited only for the first step for sequence alignment and counts matrix generation. Purchasing high-end computational power for such a small portion of a study is a massive waste of resources. One can easily overcome this need and far more effectively and efficiently process the data in the cloud. Using cloud computing provides cheap and convenient access to the necessary computing power and data sharing. The resources used can be easily customized to optimize the system for cost and time. It is also easy to process scRNA-seq data in the cloud because although the steps take a long time, the initial sequence processing steps are fixed, especially when using the single cell sequencing platform from 10X Genomics, and thus easily automated. The transformed dataset is an order of magnitude smaller which means that further analysis can be done without high-end computational resources. Importantly, these processed data are ready for analysis any downstream analysis tools; for example, an interactive free software Loupe Cell Browser from 10X Genomics is easy to use and can provide first pass analysis.

## Methods

Detailed methods for library preparation and sequencing are as described in another manuscript (Abdelfattah et al., in review).

The components for processing a scRNA-seq data was containerized and stored in the Google cloud repository. The widely used platform for generating scRNA-seq dataset is droplet based single cell capture from 10x Genomics (6). The reference implementation provides a containerized version of Cell Ranger from 10X genomics that can be used to process scRNA-seq data available in the cloud storage.

To demonstrate the versatility of this method, an online pipeline was developed with Flask (Figures 2 and 3), a comprehensive web framework in python. After data is uploaded to Google Cloud Storage (GCS), it can be submitted for post-processing by entering the URI to the data along with basic details such as a name and organism type into an online form. The framework utilizes dsub, a Google Cloud Life Sciences API, to create new VMs to process the submitted data using the Cell Ranger software (available from 10X Genomics). The web interface of a portal that captures these input variables is shown in Figure 1. To demonstrate the extensibility of this framework, we have integrated doublet removal into the pipeline, an essential next step in the scRNA-seq analysis process. Doublet detection and removal is done through Scrublet (7)., a computational framework to identify doublets in a single cell dataset. Furthermore, the initial steps of downstream analysis have been automated by incorporating Seurat, an R package to process and visualize scRNA-seq data. For example, Seurat was used in the pipeline to identify cells that likely belong to the same cell types using a clustering algorithm (Figure 4).

**Figure 1.**
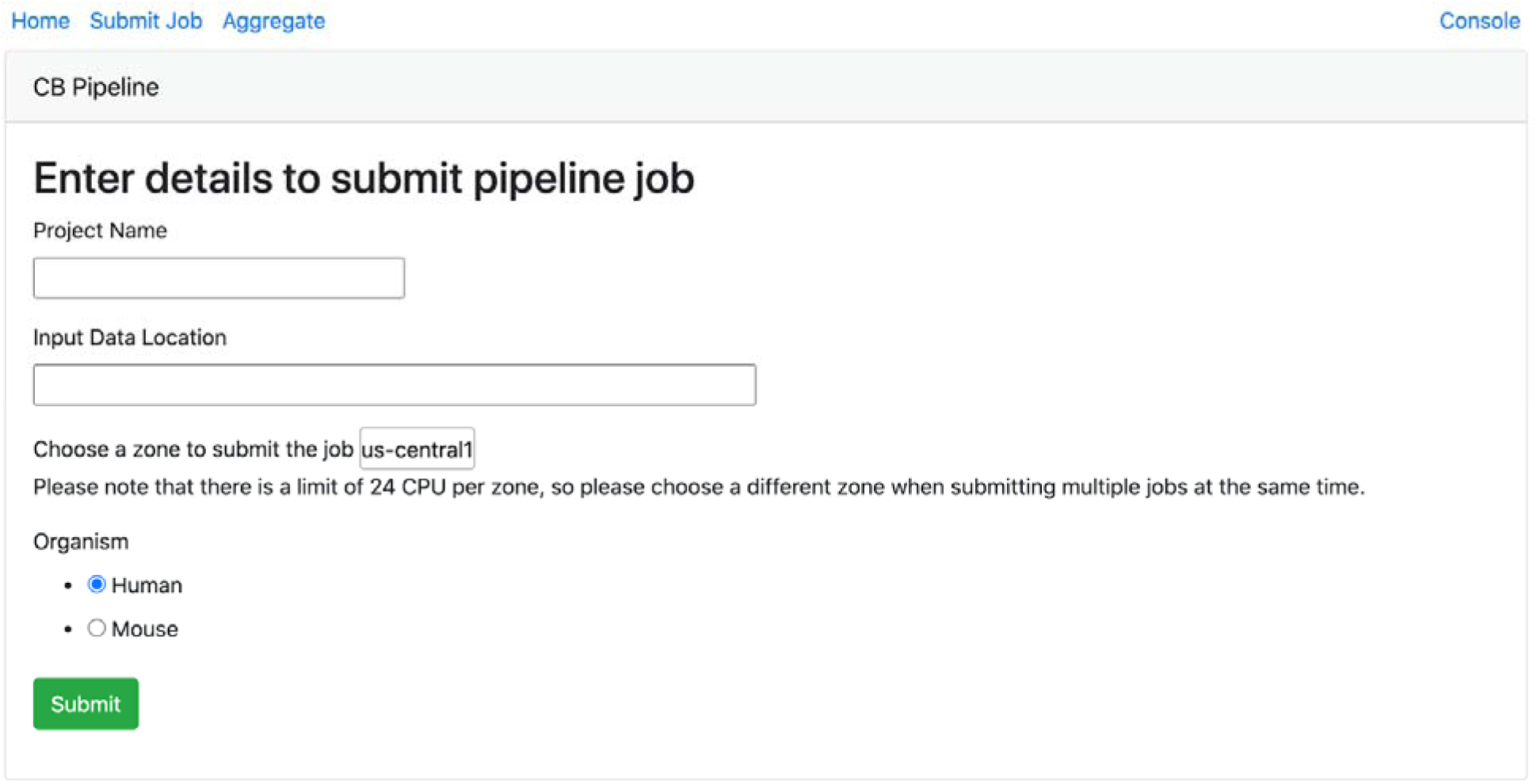
Submitting single cell data for processing is easy through the automated pipeline.

**Figure 2.**
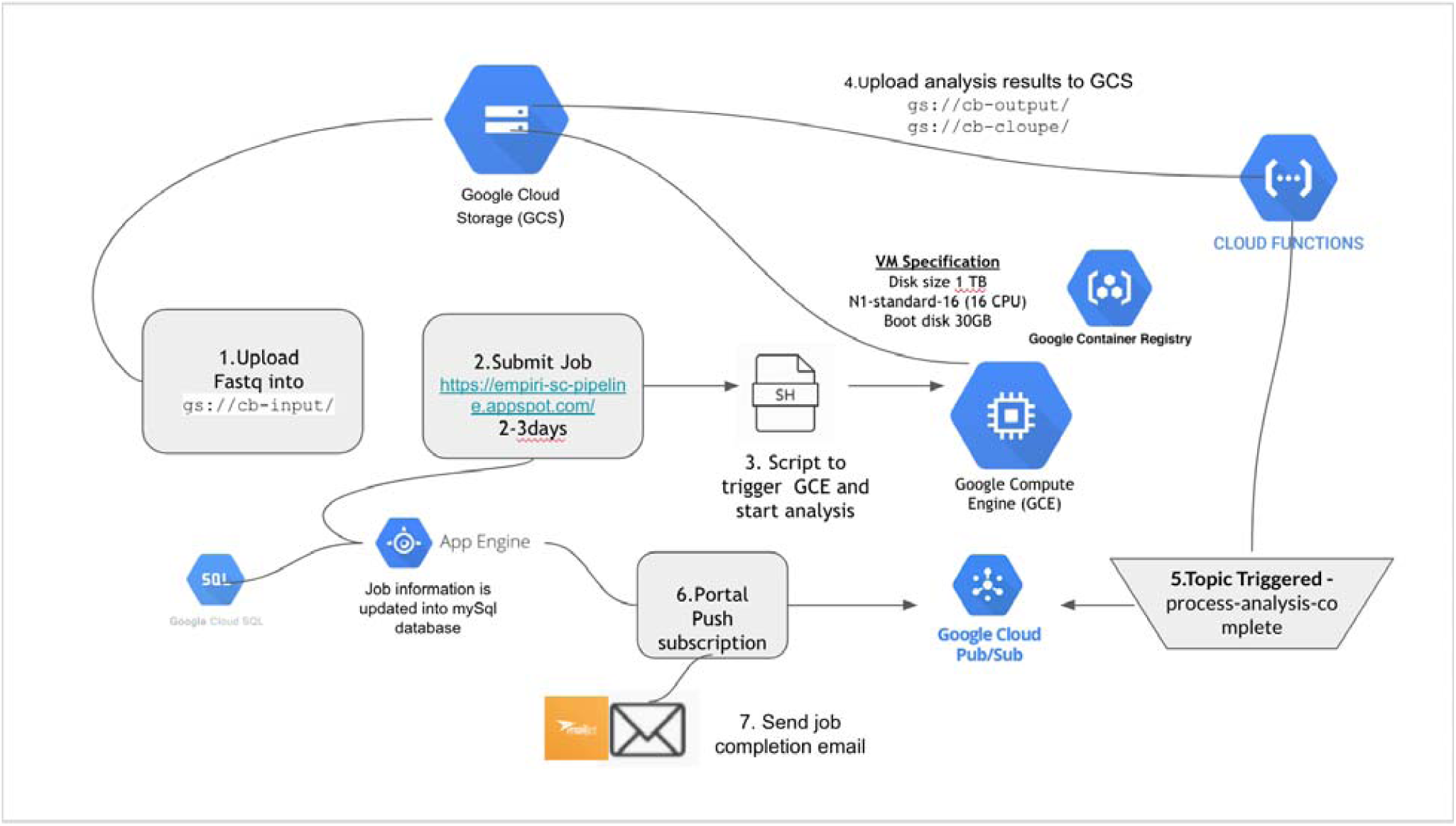
Pipeline Architecture Diagram

**Figure 3.**
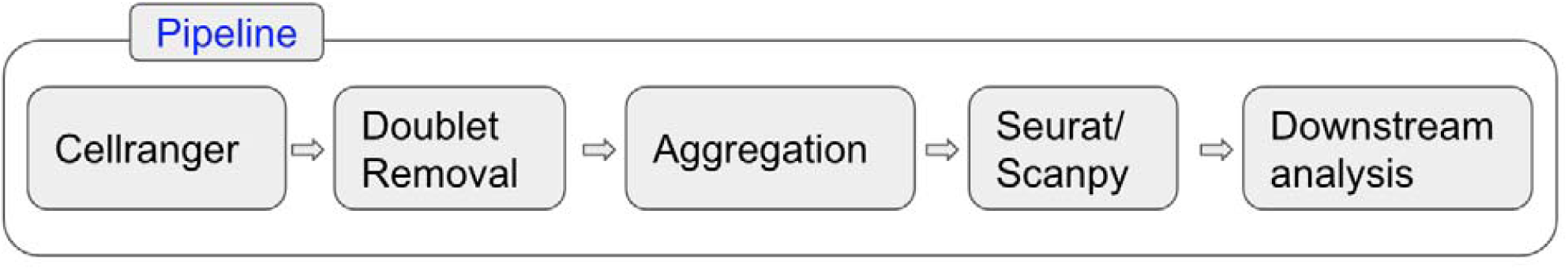
Pipeline Sample Flow Diagram

**Figure 4.**
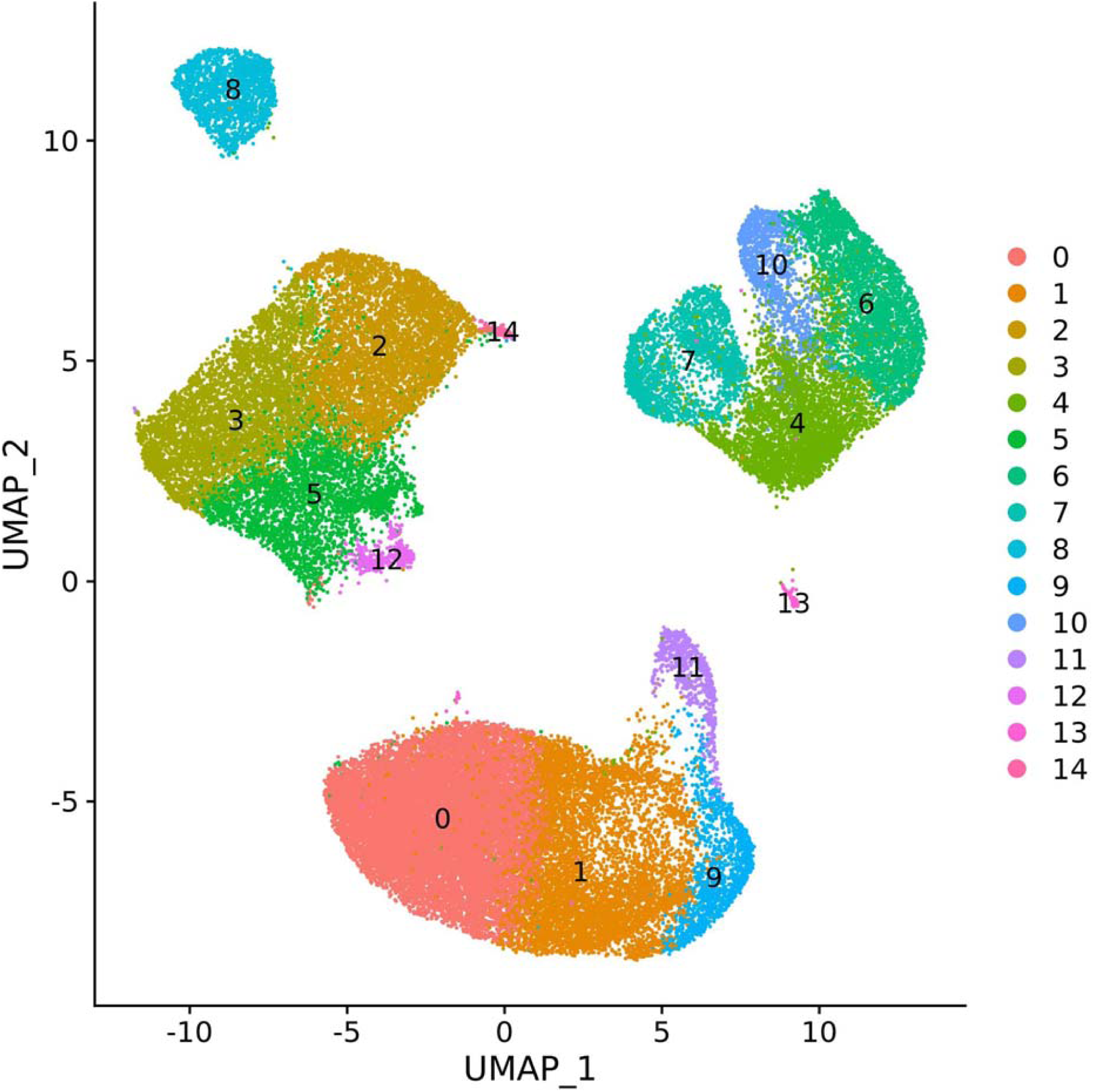
Seurat UMAP clustering of scRNA-seq data

Once sample specific information is entered, data processing will commence without any extra steps needed from the user. Deploying the web interface through the google application engine (GAE) is important to access the stored data on GCS as well as necessary cloud computing power. Computing power is obtained through the google cloud engine (GCE) which provides customizable virtual machines. Virtual machine specs are easily modified which allows the process to be optimized for cost and time. A 5000 cell PBMC dataset from 10x Genomics was used to benchmark pipeline speeds on various preset VMs from the Google Cloud Platform (Table 1). Results from this analysis revealed that processing speed is limited first by memory and then by cores, meaning that higher memory machines are more efficient. Contrary to our initial assumption, a more powerful VM is actually cheaper because it greatly reduced time for processing and lowered cost.

**Table 1.** Using a high-end machine with more memory saves on cost and time.

| Machine Type | Time (Hours) | Cost (\$) |
| --- | --- | --- |
| N1-standard-8 | 57 | 15.22 |
| N1-highcpu-32 | 19 | 15.11 |
| N1-standard-16 | 26 | 13.86 |
| N1-highmem-16 | 10 | 10.48 |
| N1-highmem-8 | 17 | 8.91 |
| N1-standard-32 | 6.75 | 7.12 |
| N1-highmem-32 | 2.5 | 5.24 |

We have developed a cloud-based framework for processing scRNA-seq data that will make scRNA-seq analysis amenable for biological research groups without rich computational resources or designated computational/bioinformatics expert. Through this pipeline, we have demonstrated how automation in the cloud is the most effective method to deal with large scRNA-seq datasets. It allows research teams to easily deal with the problem of required post-processing, allowing for greater focus on drawing biological conclusions from the data.

## Supplementary information

Supplementary data available at *Bioinformatics* online.

